# Building a genetic and epigenetic predictive model of breast cancer intrinsic subtypes using large-scale data and hierarchical structure learning

**DOI:** 10.1101/2023.06.12.544702

**Authors:** Jiemin Xie, Binyu Yang, Keyi Li, Lixin Gao, Xuemei Liu, Yunhui Xiong, Wen Chen, Li C. Xia

## Abstract

Breast cancer subtyping is a difficult clinical and scientific challenge. The prevalent Prediction Analysis of Microarray of 50 genes (PAM50) system and its Immunohistochemistry (IHC) surrogate showed significant inconsistencies. This is because of the limited training samples, highly variable molecular features and in-efficient strategies used in these classifiers. The rapid development of early screening technologies, especially in the field of circulating tumor DNA, has also challenged the subtyping of breast cancer at the DNA level. By integrating large-scale DNA-level data and using a hierarchical structure learning algorithm, we developed Unified Genetic and Epigenetic Subtyping (UGES), a new intrinsic subtype classifier. The benchmarks showed that the use of all classes of DNA alterations worked much better than single classes, and that the multi-step hierarchical learning is crucial, which improves the overall AUC score by 0.074 compared to the one-step multi-classification method. Based on these insights, the ultimate UGES was trained as a three-step classifier on 50831 DNA features of 2065 samples, including mutations, copy number aberrations, and methylations. UGES achieved overall AUC score 0.963, and greatly improved the clinical stratification of patients, as each strata’s survival difference became statistically more significant p-value=9.7e-55 (UGES) vs 2.2e-47 (PAM50). Finally, UGES identified 52 subtype-level DNA biomarkers that can be targeted in early screening technology to significantly expand the time window for precision care. The analysis code is freely available at https://github.com/labxscut/UGES.

## Introduction

Breast cancer is a malignant, complex, and highly heterogeneous tumor that overwhelmingly impacts women. Precise identification of the cancer’s subtypes is crucial to optimize treatment and outcome for patients[1]. However, breast cancer subtyping remains a difficult scientific and clinical challenge[2–4]. The most prevalent subtyping system can discriminate among four main intrinsic subtypes: Basal-like (**Basal**), Her2-enriched (**Her2**), Luminal A (**LumA**) and Luminal B (**LumB**). This system was first established in the Prediction Analysis of Microarray of 189 patients based on the expression of 50 signature genes (i.e., **PAM50** subtyping)[5]. Clinically, intrinsic subtypes were determined by a combination of surrogate Immunohistochemistry tests (i.e., **IHC** surrogate) specifically targeting estrogen receptor (ER), progesterone receptor (PR), human epidermal growth factor 2 (Her2), and Ki67 proteins[6, 7]. Given the assessment difference, it is not surprising that PAM50 subtypes and IHC surrogates demonstrated significant inconsistencies and lack of interchangeability, as proven by many large-scale multi-omics studies[8, 9]. The results, which could lead to mistreatment and misdiagnosis, call for a new identification methodology.

The inconsistencies arise from several causes. First, PAM50 subtyping is highly variable by virtue of its own nature. It is well known that mRNA expression of the same genetic information is highly variable since it tends to be disturbed by cellular, temporal and spatial noises at various stages[10–12]. Consequently, PAM50 subtyping based on the expression of only 50 genes would, almost certainly, deviate from the underlying intrinsic subtypes, as proven by many studies[13, 14]. Second, IHC surrogate to PAM50 is systematically biased. While using only four biomarkers is clinically convenient, it obviously would miss genome-wide patterns[15–17]. In fact, intrinsic subtypes are determined by a greater number of genetic and epigenetic factors, including mutation, copy number and methylation alterations, as we and others have found[18–20]. Third, current PAM50 subtyping and IHC surrogate classifications underestimated intra-tumoral heterogeneity. These classifiers were developed in the early 2000s and were trained with only a small number of samples[5, 21]. As such, they were set to miss detailed intrinsic commonalities that only become identifiable with large scale data. Our own analysis verified that PAM50 subtypes are substantially different from their IHC surrogate counterparts. Specifically, in the Molecular Taxonomy of Breast Cancer International Consortium (**METABRIC**) data, we found that 599 out of 1086 samples (55.16%) changed subtypes between PAM50 and IHC classifications (**Supplementary Table. 1**).

To address these difficulties and improve subtyping accuracy and consistency, we envision the identification of underlying genetic and epigenetic determinants of intrinsic subtypes using large-scale data and build a DNA-level subtype classifier based on these features (**Fig. 1a**). Compared to mRNA and protein changes, DNA alterations are more basic, discrete, robust, and assessable. The central dogma dictates that genetic information flows from DNA to RNA and then to protein[22]. Therefore, intrinsic subtypes of cancer cells at the DNA-level would determine downstream subtypes at expression (RNA-) and phenotype (protein-) levels. Since DNA molecules are also structurally stable[23] and not susceptible to transcriptional and translational noises, their alterations are more reliable subtyping features. Moreover, genomic and epigenomic sequencing tests of tumor biopsies and circulating tumoral materials for ascertaining DNA alterations, either targeted or genome-wide, are widely available[24]. Combining pre-screening sequencing technology advances with a DNA-level classifier could also make breast cancer subtyping possible at the early stage or even at screening, thus significantly expanding the time window for precision care.

**Fig. 1.**
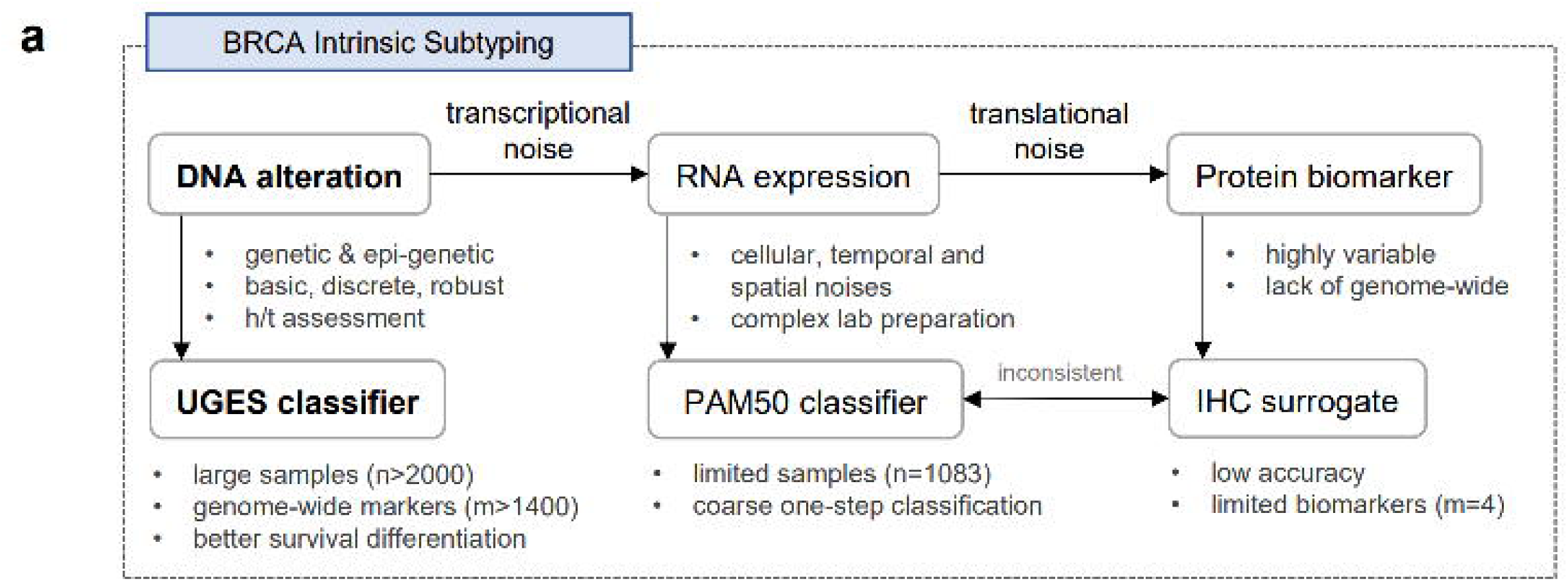
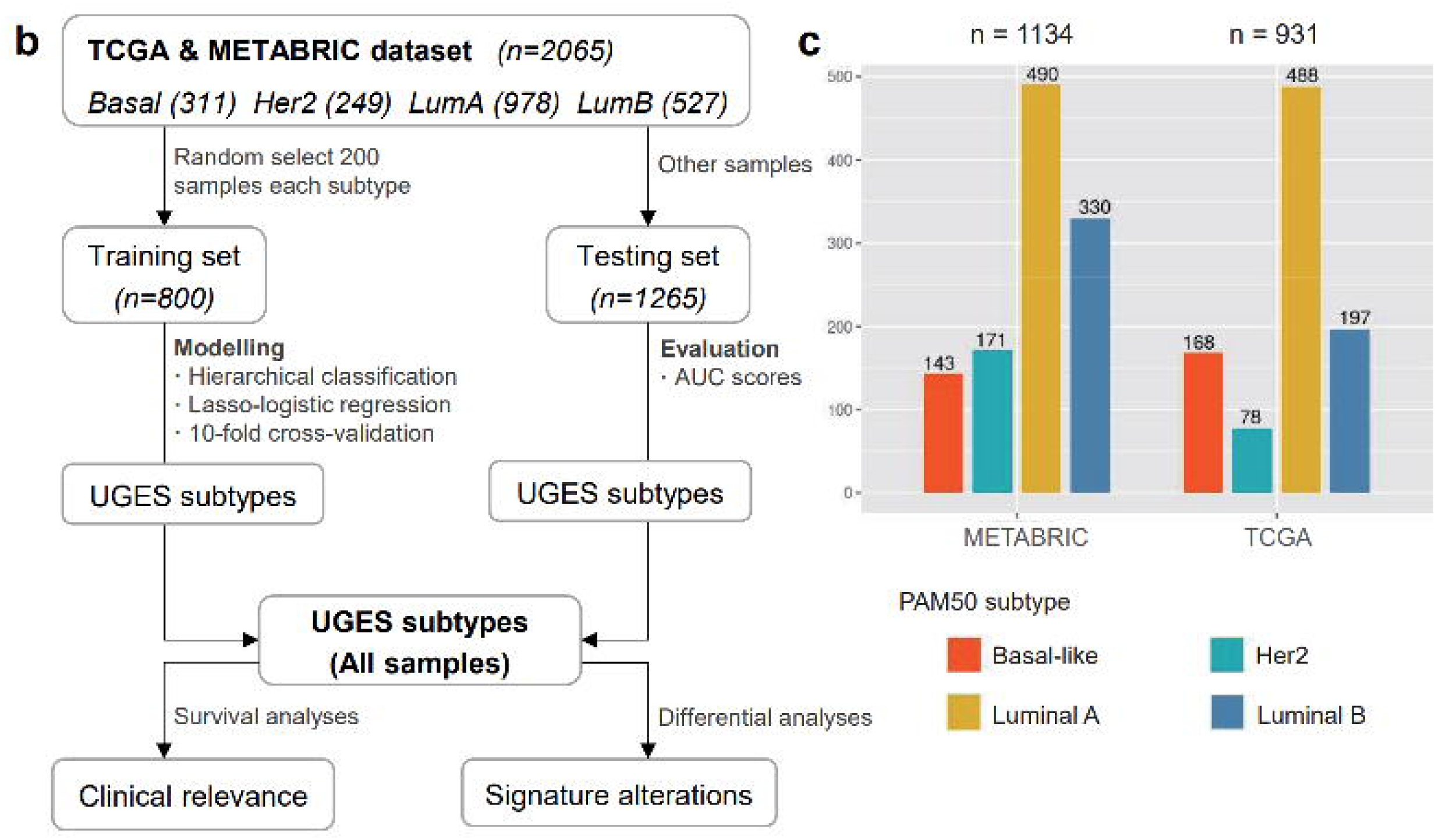

The idea that genetic and epigenetic alterations determine cancer subtypes was previously explored in breast cancer, prostate cancer, and pan-cancer studies, etc[25–27]. DNA alterations, such as mutation[28], copy number aberration (**CNA**)[29, 30] and methylation changes[31, 32], have been used entirely, or in part, for breast cancer subtyping. For example, Curtis *et al.* proposed a system of transcriptional processes depending on methylation and copy number variation and showed that CNA is a core driver of inter-tumor heterogeneity in breast cancer[33]. Islam *et al.* developed a deep neural network subtype classifier based on expression data and copy number variation data[34]. Lin *et al.* developed a deep neural networks classifier for breast cancer subtypes based on expression data, copy number variation and methylation data[35]. However, none of these studies has yet established a predictive model for determining intrinsic subtypes from DNA-level alterations alone.

By contrast, the present study leverages two large-scale (The Cancer Genome Atlas, or TCGA and METABRIC cohorts, total n=2065) (**Fig. 1c**) and multi-omics (mutation, CNA, methylation, and expression) datasets to develop a DNA-level predictive model, which we termed **UGES** (**U**nified **G**enetic and **E**pigenetic **S**ubtyping for breast cancer). Methodologically, instead of using the widely adopted one-step multi-class learning, we proposed a novel hierarchical structure learning algorithm, mimicking the clinical decision making and cancer evolution process. This change proved crucial as we will see it improves at least 7.4% accuracy over the one-step approach. Additionally, we carefully evaluated another two competitive hierarchical classification structures and demonstrated that our algorithm learned the best hierarchical classification structure, achieving the best precision-recall performance (AUC=0.963) and the best prognostic prediction power.

In UGES, we employed the Lasso-Logistic regression model as the core method for supervised label learning, which turn out an efficient approach to model high-dimensional and multi-modal data. We avoided overfitting by including the full DNA-level information and soft threshold. Based on UGES classified subtypes for the combined TCGA and METABRIC datasets, we performed survival analysis and demonstrated the clinical relevance of UGES subtypes with improved prognosis prediction over that of PAM50 system. UGES also soft-selected significant alteration features by regularizing the regression coefficients of non-informative features to zeros. Based on that and the follow-up differential analysis, we identified 52 signature-delineating DNA alterations, which collectively, sufficiently, and accurately determined the intrinsic subtype of a breast cancer.

## Materials and Methods

### Data collection and pre-processing

The Cancer Genome Atlas (**TCGA**)[36] and Molecular Taxonomy of Breast Cancer International Consortium (**METABRIC**)[33] datasets, including mutation, CNA, methylation, transcription and clinical data, were downloaded from the cBioPortal Cancer Genomics Platform (https://www.cbioportal.org)[37] and the TCGA data portal (https://portal.gdc.cancer.gov/).

We summarized all raw DNA alternations to gene level. For mutation, we mapped the downloaded mutation profiles to gene level, resulting in a 0-1 matrix, where each 𝑖, 𝑗-th cell is an indicator if the 𝑖-th gene was mutated at least once in the 𝑗-th patient. For CNA, we used the downloaded absolute copy number values, estimated by the TCGA consortium, which were already mapped to gene level by cBioPortal. For methylation, we mapped the downloaded raw methylation array profiles (HumanMethylation450 BeadChip, Illumina) to gene level, using the *FDb.InfiniumMethylation.hg19* R package. In the case of many-to-one mapping between the CpG sites and a gene, we computed and used the average methylation level for the gene.

We intersected DNA features across the two (TCGA and METABRIC) datasets and imputed missing values. We intersected and retained only methylation features available in both datasets. All DNA alteration features with more than half the values missing were excluded from the analysis. Missing values of the remaining features were imputed with 10-nearest neighbour filling, using the *impute.knn()* function of the *impute* R package[38, 39]. As a result, we included 50831 gene-level DNA features, including 16770 mutations, 25594 CNAs, and 8467 methylations.

### Lasso-Logistic regression

We applied the Lasso-Logistic regression model embedded a 10-fold cross-validation to the classification problem. The model consists of Least Absolute Shrinkage and Selection Operator (Lasso) regression and the logistic regression method, which is implemented in R, version 4.2.0, through the *glmnet* package[40]. Lasso is an efficient approach to model high-dimensional and multi-modal data, and it avoids overfitting by soft thresholding, which selects significant alteration features by regularizing the regression coefficients of non-informative features to zeros[41]. The Lasso regression model has a single hyper-parameter 𝜆 that controls the level of regularization, and we used a 10-fold cross validation on the training data to learn the optimal hyper-parameter value using the *cv.glmnet()* function. For the regularization level 𝜆, we chose the parameter value that achieves the highest classification prediction accuracy.

In order to describe this multinomial model, we used the following annotations. Suppose the response variable has 𝐾 levels 𝐺 = {1, 2, …, 𝐾}, and each sample 𝑥_𝑖_ in our study has 𝑚 = 50831 features, i.e., 𝑥_𝑖_ = {𝑥_𝑖,1_, 𝑥_𝑖,2_, …, 𝑥_𝑖,𝑚_}, then we model:

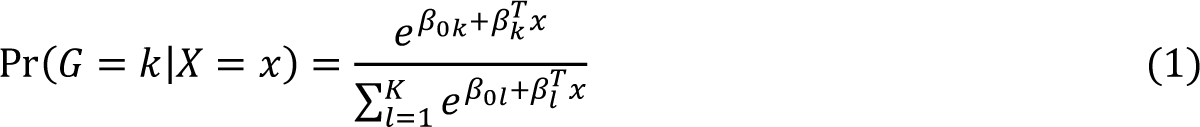

Let 𝑌 be the 𝑁 × 𝐾 indicator response matrix, with elements 𝑦_𝑖𝑙_ = 𝐼(𝑔_𝑖_ = 𝑙). Then the Lasso elastic net penalized negative log-likelihood function becomes:

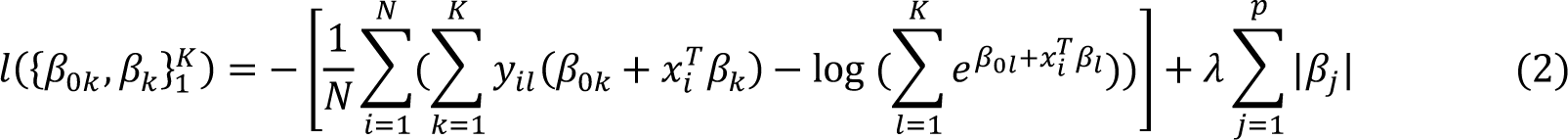

where 𝛽 is a 𝑝 × 𝐾 matrix of coefficients, 𝛽_𝑘_ refers to the 𝑘th column (for outcome category 𝑘), and 𝛽_𝑗_ the 𝑗th row (vector of 𝐾 coefficients for variable 𝑗), 𝜆 is the penalty parameter. In our study, 𝐾 = 4, 𝑁 = 2065.

### Hierarchical structure learning algorithm

To construct the optimal classification structure, we extended the tree structure learning algorithm proposed by Bengio *et al.*[42] and applied this image classification algorithm to our data. In general, the first step of our algorithm is to find a metric that can represent the similarity relationship between classes, and the second step is to construct a clustering structure based on this metric. The algorithm flow is shown in **Algorithm 1**.

**Algorithm 1:**
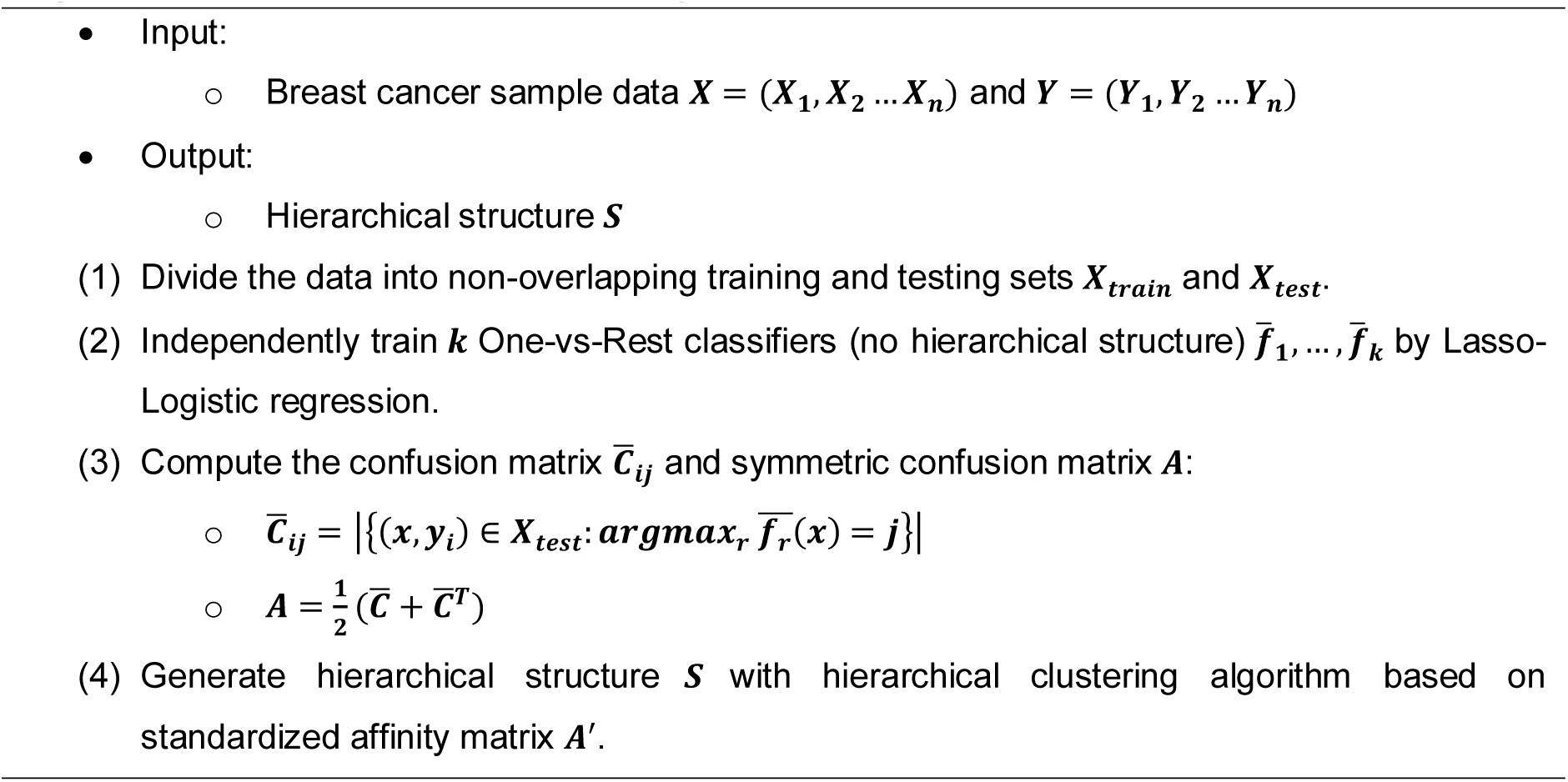
Hierarchical structure learning

In the algorithm used in this paper, the similarity measure between different classes is obtained by training 𝑘 One-vs-Rest classifiers to obtain the confusion matrix ^𝑪̅^_𝒊𝒋_ ∈ 𝑹^𝑲×𝑲^. If category 𝑖 is more similar to 𝑗, for sample 𝑥_𝒊_, it is more likely to be classified as a positive case in the classifier 𝑓_𝑗_, i.e., 𝑓_𝑗_ (𝑥_𝑖_) = 𝑗. We convert this confusion matrix into an inter-class affinity matrix, which is a symmetric matrix that represents the similarity or correlation between each pair of classes. Each element of the affinity matrix represents the degree of affinity or correlation between two classes, for example, the (𝑖, 𝑗)-th element of the matrix represents the degree of similarity or correlation between class 𝑖 and class 𝑗. For the choice of clustering algorithm, we used the hierarchical clustering algorithm, which more directly models the affinity between classes. First, each class is considered as a separate class, and then the two classes with the highest similarity are combined into a new class, until finally only one class remains.

### Relative importance

Genetic and epigenetic determinants of intrinsic subtypes were identified by the Lasso-Logistic regression model, allowing for the quantification of effects for three DNA alteration classes. Since Lasso-Logistic regression allows for high-dimensional variable compression and selection, the Lasso-selected alterations and their corresponding regression coefficients were summarized to assess their importance. In total, 236, 367 and 530 DNA alterations were selected from the *B-vs-(H,LA,LB)*, *H-vs-(LA,LB)* and *LA-vs-LB* sub-classifiers of UGES, respectively (**Supplementary Table. 2**).

We applied relative importance to measure the total effect of each DNA alteration class[43–45]. An easy and common method for estimating the relative importance of a DNA alteration class is to express the sum of each regression coefficient in this class as a percentage of the sum of all coefficients. Suppose 𝑓 = {𝑉, 𝐶, 𝑀} is the set of three DNA alteration classes, where 𝑉 denotes the set of all mutation alterations, 𝐶 denotes CNA alterations, and 𝑀 denotes methylation alterations; then, the relative importance 𝑅𝐼_𝑖_ of type 𝑖 can be calculated by equation (3):

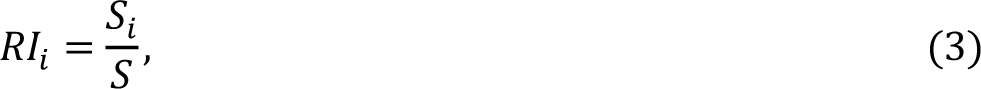

where 𝑆_𝑖_ = ∑_𝑖 ∈ 𝑓_|𝑐_𝑖_ | is the sum of all absolute regression coefficients for a certain DNA alteration class and 𝑆 = ∑_𝑓_ 𝑆_𝑓_.

### Survival analysis

Survival analyses were performed in R, version 4.2.0, using the *survival* and *survminer* package with Overall Survival (OS) as the primary endpoint. In univariate analysis, survival situations were compared using the Log-rank test.

In multivariate analysis, patient age was included as a covariate. Cox regression (or Cox proportional hazards regression) was used to analyze the effect of UGES subtypes and age, using the *coxph()* R function. The forest plot showed survival outcomes according to different UGES subtypes and age, using a resampled balanced patient dataset (304 samples for each subtype) as reference. Analyses involving multiple comparisons were adjusted using the Benjamini-Hochberg (BH) FDR method, which controls the False Discovery Rate (FDR) with the Holm–Bonferroni method for multiple hypothesis testing. All statistical tests were determined to be significant based on the BH-corrected significance level of 0.05.

### Differential analyses

Differential analyses were conducted in R, version 4.2.0, on 112 DNA alteration features, including all top 10% alterations in each sub-classifier by coefficient values. For mutation data, we applied the Chi-Square Test using *chisq.test()* R functions. For copy number aberration data, we applied the Kruskal-Wallis Test conducted by *kruskal.test()* R function. For methylation data, we applied the Welch ANOVA Test using the *oneway.test()* R function and set parameter *var.equal = FALSE*. Features were considered significally different if the BH-corrected test p-value was < 0.01. All tests were two-sided.

## Results

### Cohort characteristics

We downloaded and analyzed DNA-level and clinical data for a total of 2065 samples, including 931 TCGA samples and 1134 METABRIC samples (**Fig. 1c**). We assigned these samples to three breast cancer cohorts: the TCGA only cohort, the METABRIC only cohort, and the combined TCGA+METABRIC cohort. The descriptive statistics of demographic (age) and relevant clinical variables (ER status, PR status, Her2 status) were presented for each cohort, as shown in **Table. 1**.

Analyses of clinical variables also yielded confirmatory statistics suggesting that the cohorts were representative of the general breast cancer population, and thus suitable for our integrative analysis. Elder patients had a statistically significant worse survival rate as expected[46, 47], with a relative increase of death risk of approximately 4% (HR = 1.039, 95% CI = [1.029, 1.049], p-value < 2e-16, on average), demonstrating that the clinical variables were generally consistent across the cohorts. Patients with positive ER status or negative Her2 status also had a statistically significant improved survival as expected[48, 49], with a relative reduction of death risk by >25% or >64% (positive ER status: HR = 0.749, 95% CI = [0.563, 0.997], p-value = 0.0474; positive Her2 status: HR = 1.643, 95% CI = [1.244, 2.169], p-value = 0.0005).

### UGES -- a DNA-level classifier sufficiently and accurately identifies intrinsic subtypes

We first built a new DNA-level intrinsic subtype classifier. One drawback of multi-omics subtyping is its mixed and undifferentiated use of molecular features, whereas a restrained-yet-effective DNA-level classifier will improve subtyping interpretability, applicability, and robustness. Yet, we have to determine which classification strategy and what DNA-level features to use in such a classifier. For both the TCGA and METABRIC datasets, the DNA-level alterations including mutations, CNAs and methylations are fully available. We first used this full DNA information to evaluate candidate hierarchical classification strategies. We then evaluated the informativeness of individual DNA-level alteration categories by applying all potential classification strategies.

We obtained a three-step hierarchical classification structure for breast cancer intrinsic subtypes based on the proposed hierarchical structure learning algorithm (see **Algorithm 1 in Materials and Methods**), and simply termed it **(B,(H,(LA,LB)))** by the Newick notation (**Fig. 2a**)[50]. In the notation, B, H, LA and LB denote the **B**asal, **H**er2, **L**um-**A** and **-B** subtypes respectively. In the **(B,(H,(LA,LB)))** strategy, Basal and other subtypes were grouped at the first classification step, then Her2 subtypes was separated, and finally Luminal-like subtypes were grouped.

**Fig. 2.**
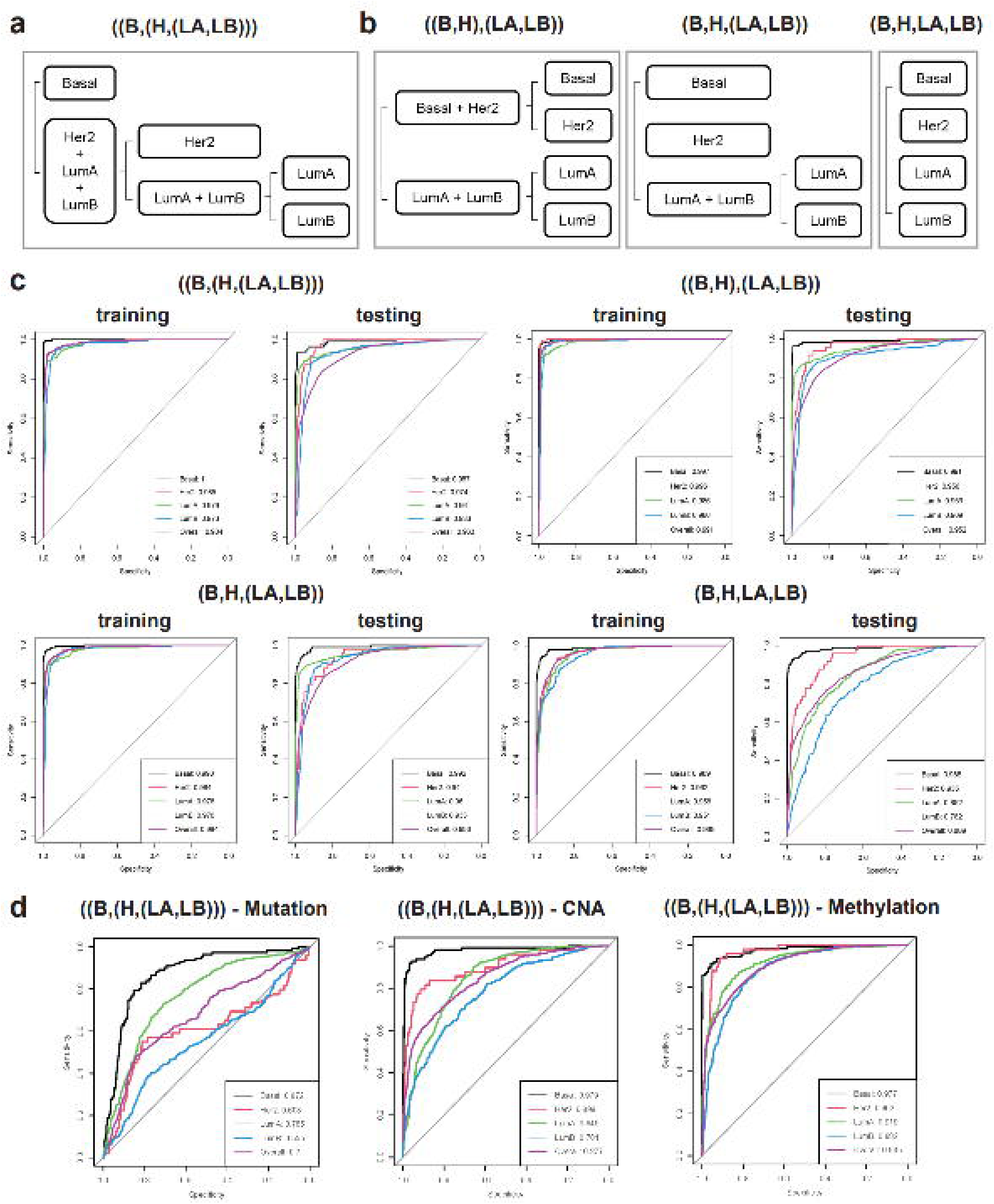

We also examined three competitive hierarchical classification strategies (**Fig. 2b**). The first and most commonly used strategy is simple one-step multi-class learning[51], or **(B,H,LA,LB)**, which classifies 4 intrinsic subtypes at once. In the **(B,H,LA,LB)** subtype grouping, all included subtypes are classified in the same single step. Another strategy we included is **(B,H,(LA,LB))** two-step classification, which arises from the clinical decision process in which a patient was first tested for Basal and Her2 biomarkers and then tested for LumA and LumB sub-markers for subtyping[51, 52]. (B,H,(LA,LB)) is reasonable because the gene expression of Luminal-like subtypes is similar[53, 54], as compared with other intrinsic subtypes. Finally, given the under-presence of Basal and Her2 subtypes in the discovery cohorts, we proposed a new strategy **((B,H),(LA,LB))**, where Basal and Her2 subtypes were grouped at the first classification step to boost their sample number, in order to balance training samples and increase label learning efficiency and generalizability[55].

To compare strategies, we divided the combined TCGA+METABRIC samples into non-overlapping training and testing subsets (**Fig. 1b**). We trained and cross-validated DNA-level classifiers based on all three strategies with the training subset, and then applied them to the testing subset. We used Receiver Operating Characteristic (**ROC**) curves and Area Under Curve (**AUC**) scores as performance evaluation metrics. For technical details, please refer to **Materials and Methods**.

We found that the (B,(H,(LA,LB))) strategy generated by our hierarchical structure learning algorithm best identifies intrinsic subtypes. In fact, all the multi-step strategies had high performance and were virtually indifferentiable overall. As shown in **Fig. 2c**, the training and testing overall AUC scores (averaged over all subtypes) of (B,(H,(LA,LB))) were 0.984 and 0.963, respectively, while those for ((B,H),(LA,LB)) were 0.991 and 0.952, for (B,H,(LA,LB)) were 0.984 and 0.956, and for (B,H,LA,LB), they were only 0.965 and 0.889. The fact that AUC scores for these strategies were greater than 0.9 supports our reasoning that DNA-level information is sufficient and robust for identifying intrinsic subtypes accurately.

All multi-step strategies grouped Luminal-like subtypes at the first step, resulting in a substantial improvement by increasing the overall AUC by at least 6.3% over the one-step multi-class strategy, as adopted by PAM50 subtyping and many other multi-omics subtyping methods. This result gave evidence of a common genetic and epigenetic basis for Luminal-like tumors, as they diverged further from Basal and Her2 subtypes. Such commonality was not identifiable using a coarse one-step multi-class approach. We also compared the three strategies using the ROC curves of the combined set by the one-vs-one method (**Supplementary Fig. 1**). It was obvious that (B,H,LA,LB) performed the worst at distinguishing LumA and LumB (AUC = 0.828) compared to the three-step classifier (AUC=0.948 in (B,(H,(LA,LB)))) and two-step classifiers (AUC=0.941 in ((B,H),(LA,LB)) and 0.951 in (B,H,(LA,LB))), improves the separation accuracy by 11.3% at distinguishing Luminal A and B subtypes, the most difficult separation.

We found the (B,(H,(LA,LB))) strategy to be the best for identifying Her2 and Luminal-like subtypes. In terms of the testing set, its AUC scores are 0.987, 0.974, 0.96 and 0.933 for Basal, Her2, LumA and LumB subtypes, respectively, while those for ((B,H),(LA,LB)) are 0.991, 0.956, 0.953 and 0.909, for (B,H,(LA,LB)) are 0.992, 0.94, 0.96 and 0.933, and for (B,H,LA,LB) are 0.988, 0.935, 0.852 and 0.782, respectively (**Fig. 2c**). In fact, (B,(H,(LA,LB))) gained considerably (AUC +0.018) in identifying the Her2 subtype, performed best in identifying LumA and LumB subtypes, and remained most competitive with only a slight loss in identifying Basal cases (AUC -0.005). Considering the best performance of (B,(H,(LA,LB))), which was also generated by the hierarchical structure learning algorithm, we selected it as the ultimate classification strategy to implement in the UGES classifier.

We additionally found that classifiers using all DNA-alterations generally worked much better than those using only single feature. For example, using the testing set, the all-feature (B,(H,(LA,LB))) classifier had an overall AUC=0.963, while the AUCs for mutation-, CNA- and methylation-only classifiers were 0.7, 0.877 and 0.935 respectively, which are all lower. Subtype-wise, the all-feature (B,(H,(LA,LB))) classifiers also had consistently better performance than the corresponding single-feature classifiers (**Fig. 2d**).

The same pattern is true for classifiers using the other three strategies in that the all-feature classifiers almost all had higher overall and subtype-wise AUC scores than single-feature classifiers (**Supplementary Fig. 2**), except for the methylation-only (B,H,LA,LB) classifier, which performed better (AUC = 0.943) than the all-feature classifier (AUC = 0.935) in identifying the Her2 subtype. This could be caused by a model overfitting problem specific to the methylation training data, as we discussed later (see **Discussion**). Based on these findings, we decided to include all mutation, CNA and methylation features into building the UGES classifier and finally implemented it as a two-step all-feature (B,(H,(LA,LB))) classifier with the best overall performance (AUC=0.963) in this benchmark.

### DNA methylation has the main effect in defining subtypes

We studied the **relative importance** of each DNA feature based on UGES defined intrinsic subtypes. Biologically, genetic and epigenetic alterations can have different effects on intrinsic subtypes via varied biological mechanisms[56, 57], including gene functional disruption, gene dosage change, and genome accessibility change. The availability of full DNA-level data on a large-scale allowed us to quantify these effects based on UGES subtyping, which could help in exploring subtype-wise early detection and precise treatment options. Relative importance is a metric calculating the relative contribution of predictive variables (see Eq.**(3) in Materials and Methods**) to the response. The relative importance of DNA features in the UGES sub-classifiers *B-vs-(H,LA,LB)*, *H-vs-(LA,LB)*, and *LA-vs-LB* were shown in **Table. 2**.

We found that DNA methylation change was consistently the most important alteration in defining all sub-classifiers. The relative importance of methylation features was 53.97%, 55.08% and 51.62% for *B-vs-(H,LA,LB)*, *H-vs-(LA,LB)*, and *LA-vs-LB* sub-classifiers, respectively, all over 50%. In general, CNA is a remote second in importance, which accounted for 33.35%, 25.15% and 19.76% of the total importance in the respective models, and mutation is in third place, accounting for 12.68%, 19.78%, and 28.63% of the total in the respective models. Consistent with our quantitative assessment here, the methylation-feature-only classifiers had the highest AUC scores for all subtypes (see **Fig. 1d**), suggesting that methylation change as a whole is probably the most prominent feature in defining the intrinsic status of cancerous cells, which were also reported in previous studies alphabet in smaller sample sizes[58, 59].

Either CNA or mutation features were the second most important depending on the sub-classifier. CNA features had the second largest relative importance in the *B-vs-(H,LA,LB)* and *H-vs-(LA,LB)* sub-classifiers, 33.35% and 25.15%, respectively, but the smallest relative importance in the *LA-vs-LB* sub-classifier, at 19.76%. Mutation features had the second largest relative importance in the *LA-vs-LB* sub-classifier at 28.63% but the smallest in the *B-vs-(H,LA,LB)* and *H-vs-(LA,LB)* sub-classifiers, at 12.68% and 19.78%, respectively. Notably, the relative importance of mutation features was the second highest (28.63%) in the *LA-vs-LB* sub-classifier, but the lowest in the other sub-classifiers, suggesting that mutations were crucial in distinguishing LumA and LumB subtypes, as reported by previous research[60].

### UGES defined intrinsic subtypes show improved clinical relevance

We found that UGES subtypes were substantially different from PAM50 subtypes. The alluvial plot and the confusion matrix showed that subtype reassignment occurred for 11.57% (239 in 2065 samples) (**Fig. 3a, b**). The reassignment better resolve LumA and LumB subtypes classification. Among Basal, Her2, LumA and LumB PAM50 subtypes, 5.79%, 7.63%, 15.64% and 9.87% changed subtypes by UGES, with the most frequent change being 11.25% LumA (PAM50) to LumB (UGES). This is plausible because Luminal-like types are genetically and epigenetically more similar compared to others and are the most challenging to resolve.

**Fig. 3.**
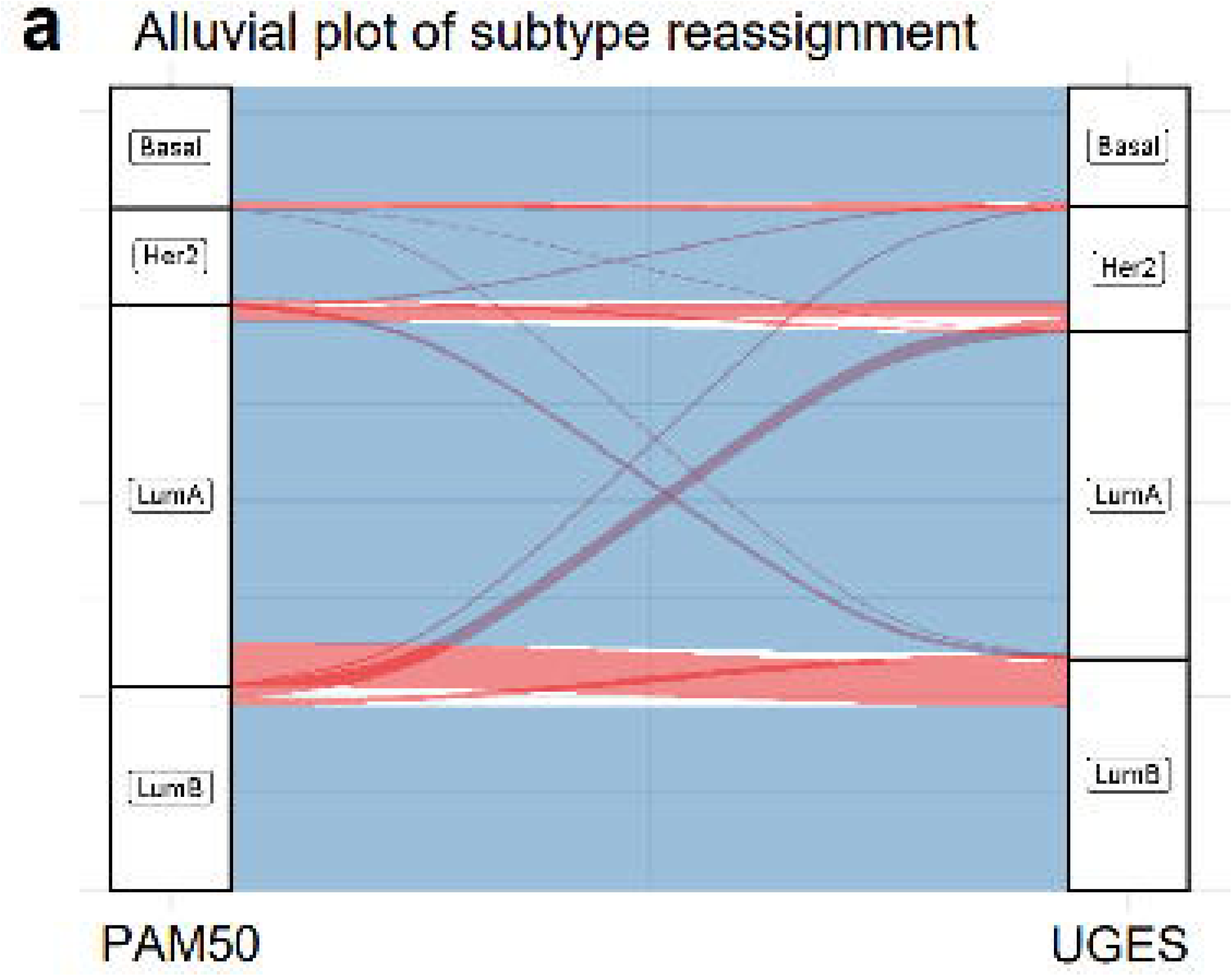
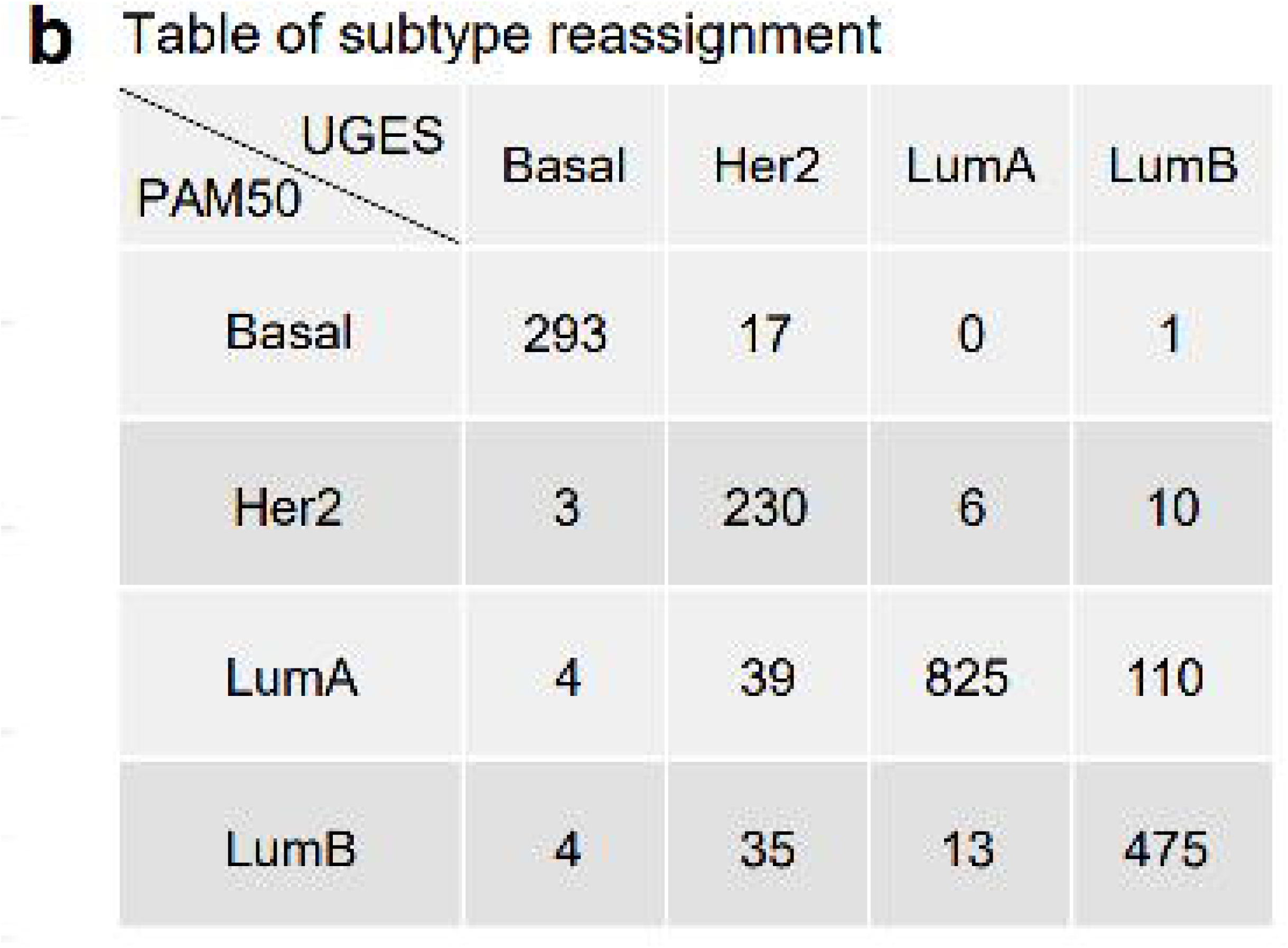
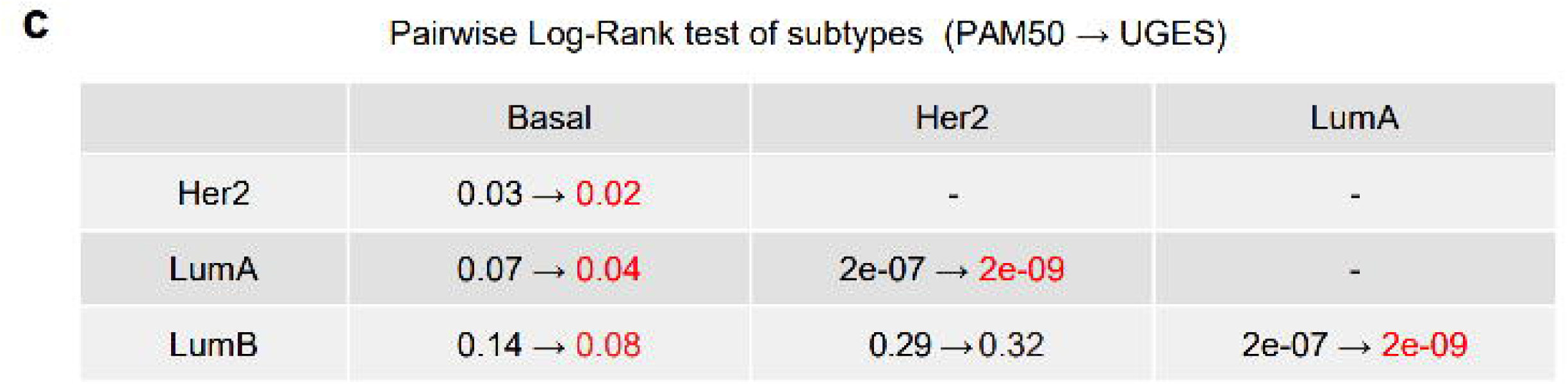
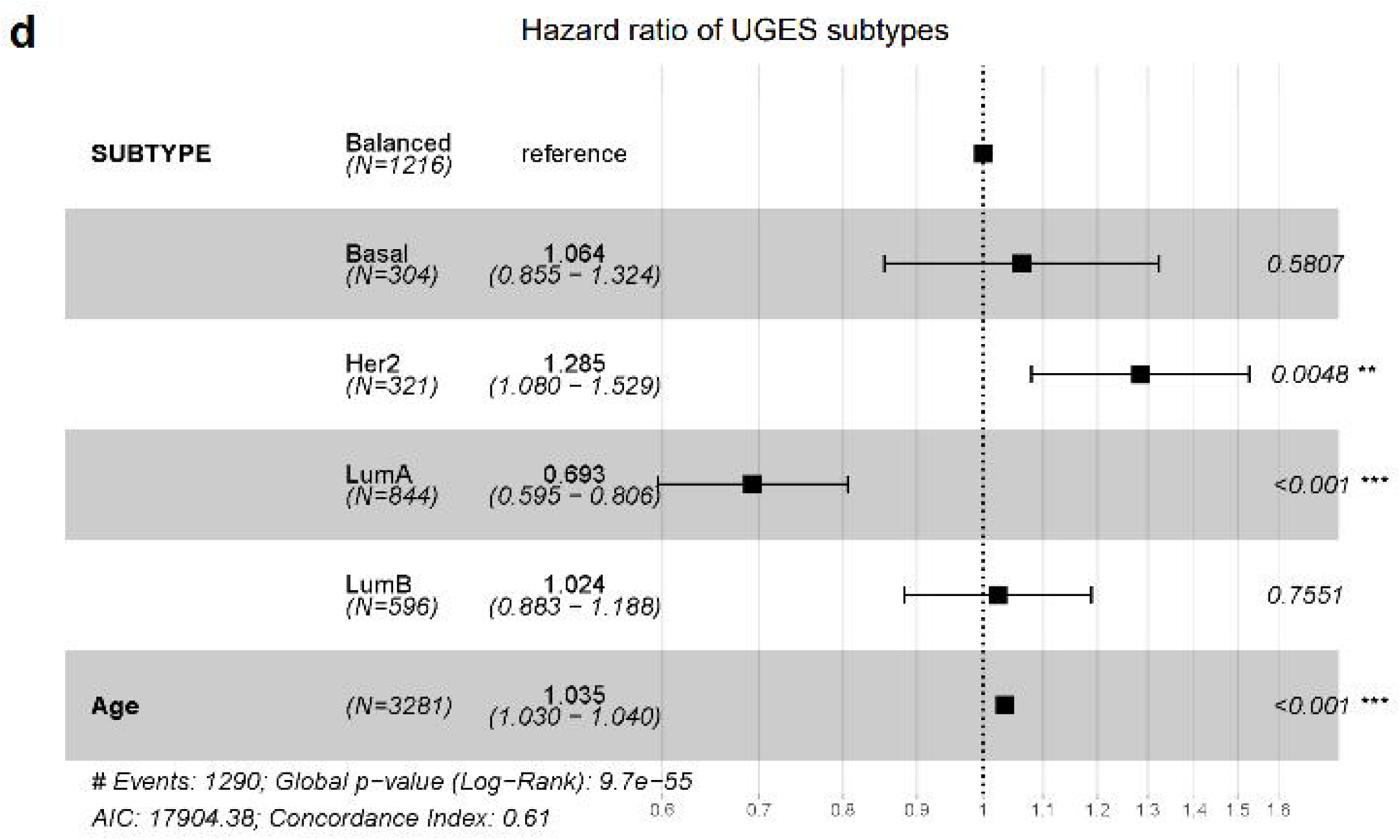

UGES increased the model sensitivity to detect the most difficult Her2 intrinsic subtype (+28.92% after reassignment). 94.44% of Basal (PAM50) subtype changed to Her2 (UGES) subtype (17 out of 18 switching) and 67.31% of LumB (PAM50) subtype changed to Her2 (UGES) subtype (35 out of 52 switching), while the frequency of change to other subtypes was much lower, with 0% and 5.56% of Basal (PAM50) subtype changed to LumA and LumB (UGES) subtypes, and 7.69% and 25% of LumB (PAM50) subtype changed to Basal and LumA (UGES) subtypes, respectively.

We then demonstrated that UGES subtypes were more predictive of patients’ overall survival as evidence of its validity and clinical relevance. We found that UGES subtypes showed improved prognostic value with better overall survival (OS) stratification compared to PAM50 subtypes (**Fig. 3c, d**). The results were based on both univariate and multivariate survival analyses (see **Materials and Methods**). Based on univariate survival analysis, in 5 out of 6 pairwise comparisons (each comparing two subtypes), we found that the survival of UGES subtypes was more distinguishable from each other than PAM50 subtypes (**Fig. 3c**). For example, the statistical significance of survival differences among these UGES subtypes were increased (smaller p-values), except between Her2 and LumB subtypes, proving that the UGES subtypes are more clinically relevant. Moreover, under UGES, the survival differences between the LumA and Basal subtypes became statistically significant (p-value decreased from 0.07 to 0.04). In general, UGES subtyping resulted in sharper difference between clinical intrinsic subtype groups.

In multivariate survival analysis, where age was added as a risk factor, UGES subtyping also showed improved clinical relevance. Using a resampled balanced sample set as a benchmark, UGES subtypes were also found to better stratify the intrinsic subtypes with regard to their survival difference (p-value = 9.7e-55), as compared to PAM50 subtypes (p-value = 2.2e-47) (**Fig. 3d**). For example, the significance of the differences between Her2 and the other subtypes had increased, changing from statistically insignificant to significant (p-value decreased from 0.0557 to 0.0048). Moreover, LumA subtype had higher overall survival rate (hazard ratio < 1) compared to the other subtypes, verifying the better prognosis of LumA patients, as reported by others[61]. These findings proved that UGES defined subtypes yielded more clinically distinguishable patient groups, which will be useful for more precise clinical management.

### UGES identified signature subtype-delineating alterations

Unravelling genetic and epigenetic heterogeneity and identifying subtype-specific markers of breast cancer are the keys to developing targeted therapies that will lead to improved outcomes[62]. Here, we conducted differential analyses using 112 genetic and epigenetic features selected as the top 10% most important features in each UGES sub-classifier (see **Materials and Methods**). As a result, we identified 52 subtype-delineating signature alterations. These signature DNA features were presented in **Table. 3** regarding their corresponding subtypes. Heatmap plots of signature DNA alteration profiles in **Fig. 4** also showed patterns consistent with the results given in **Table. 3**.

**Fig. 4.**
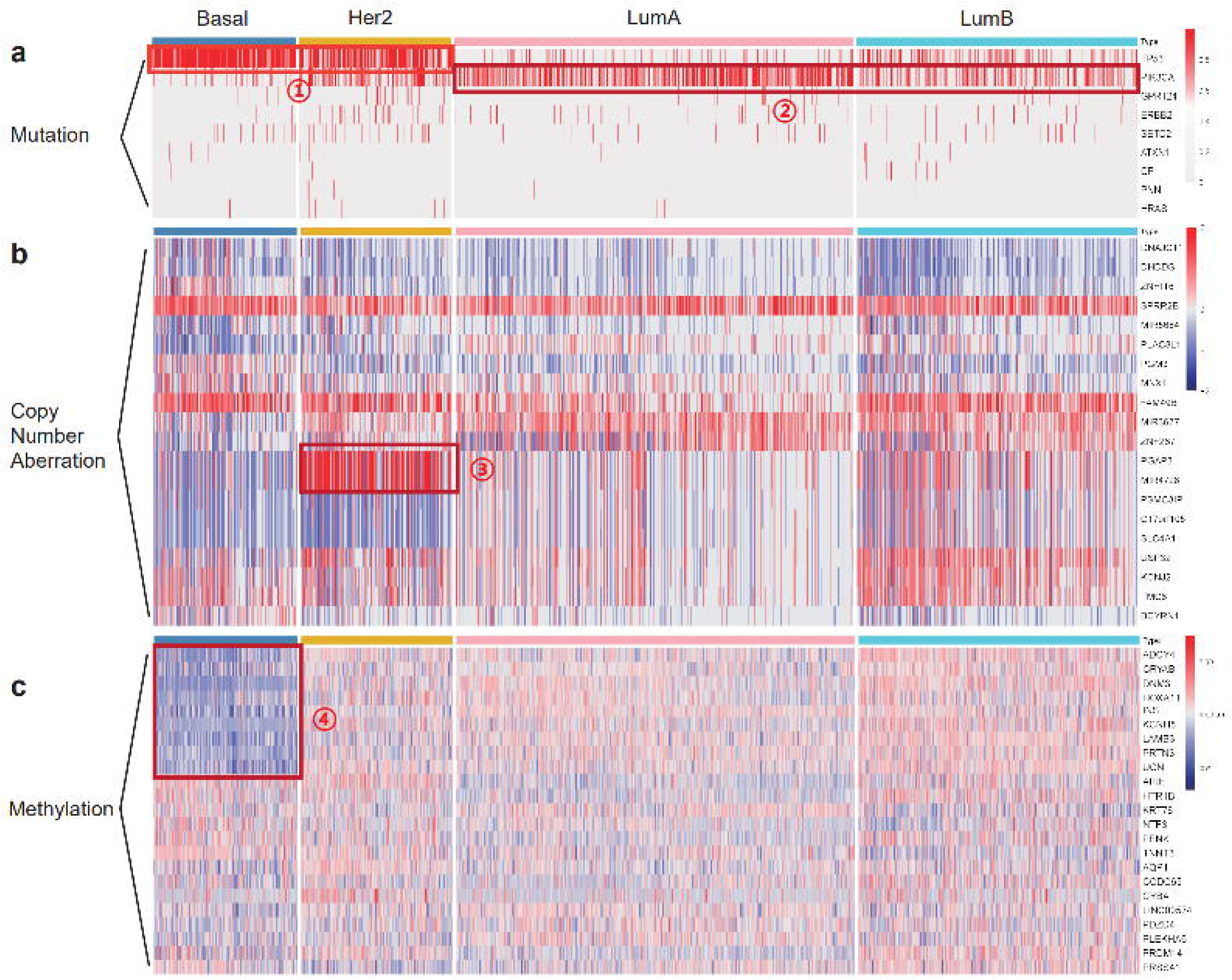

**Fig. 5.**
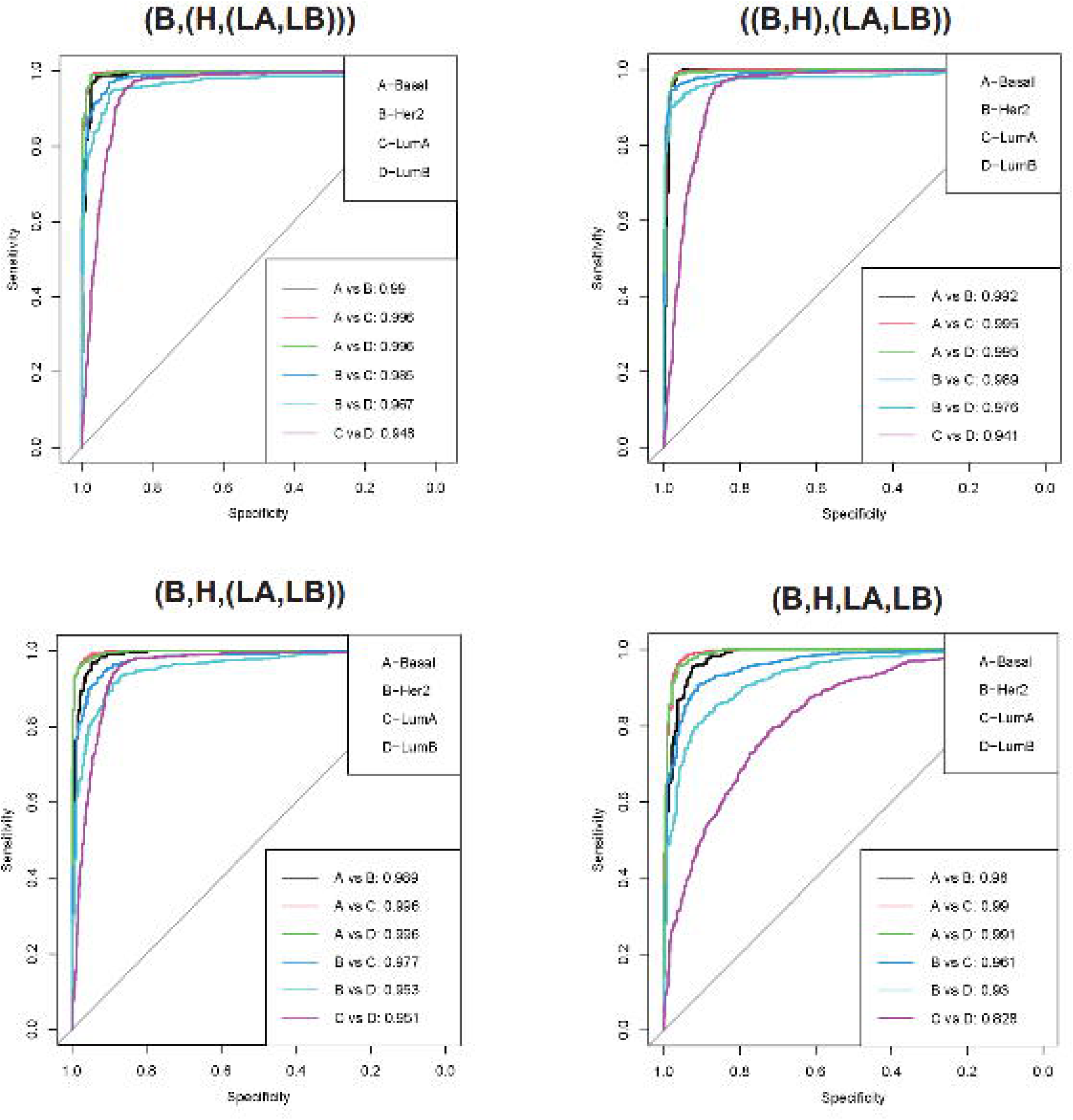
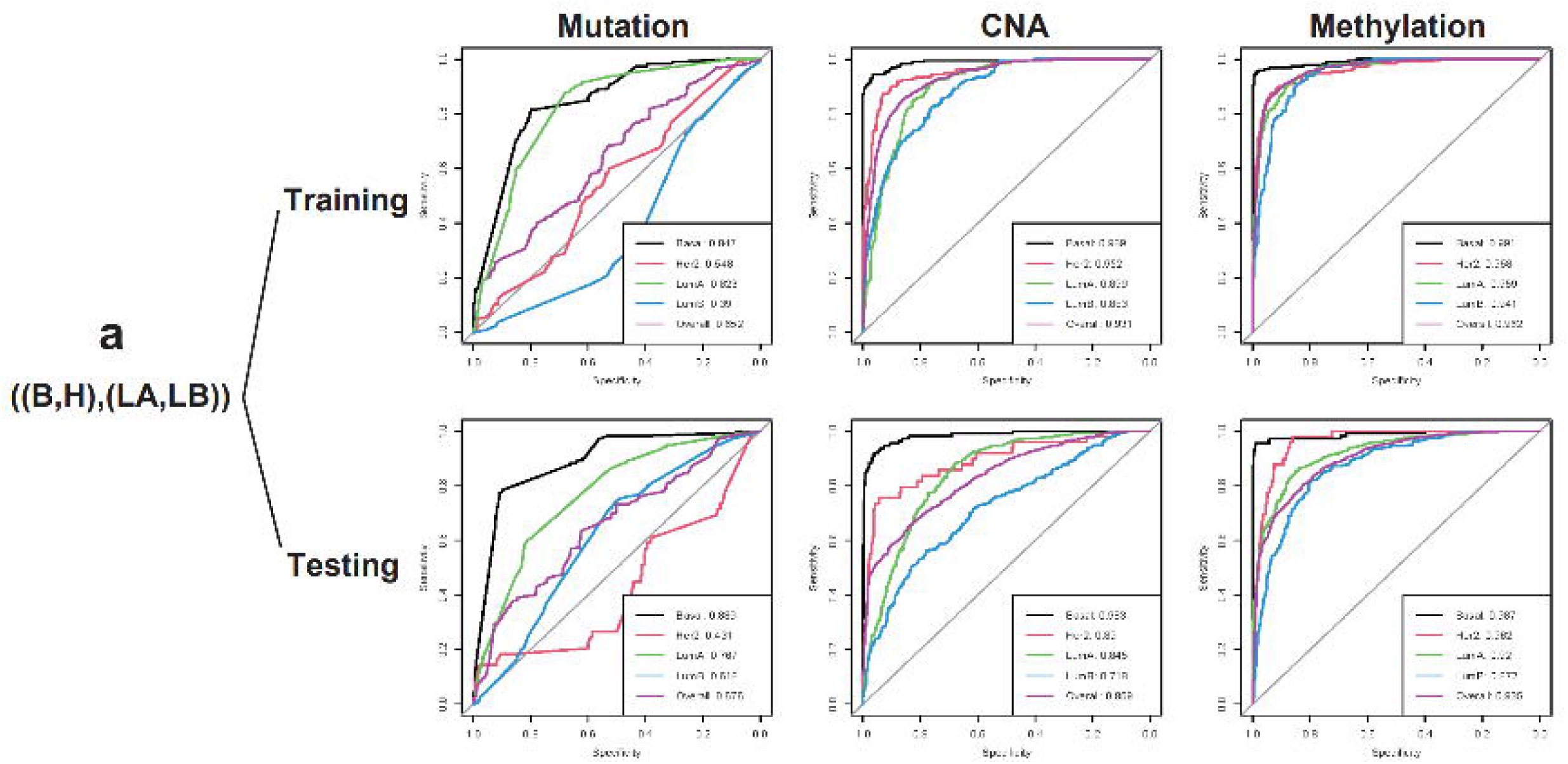
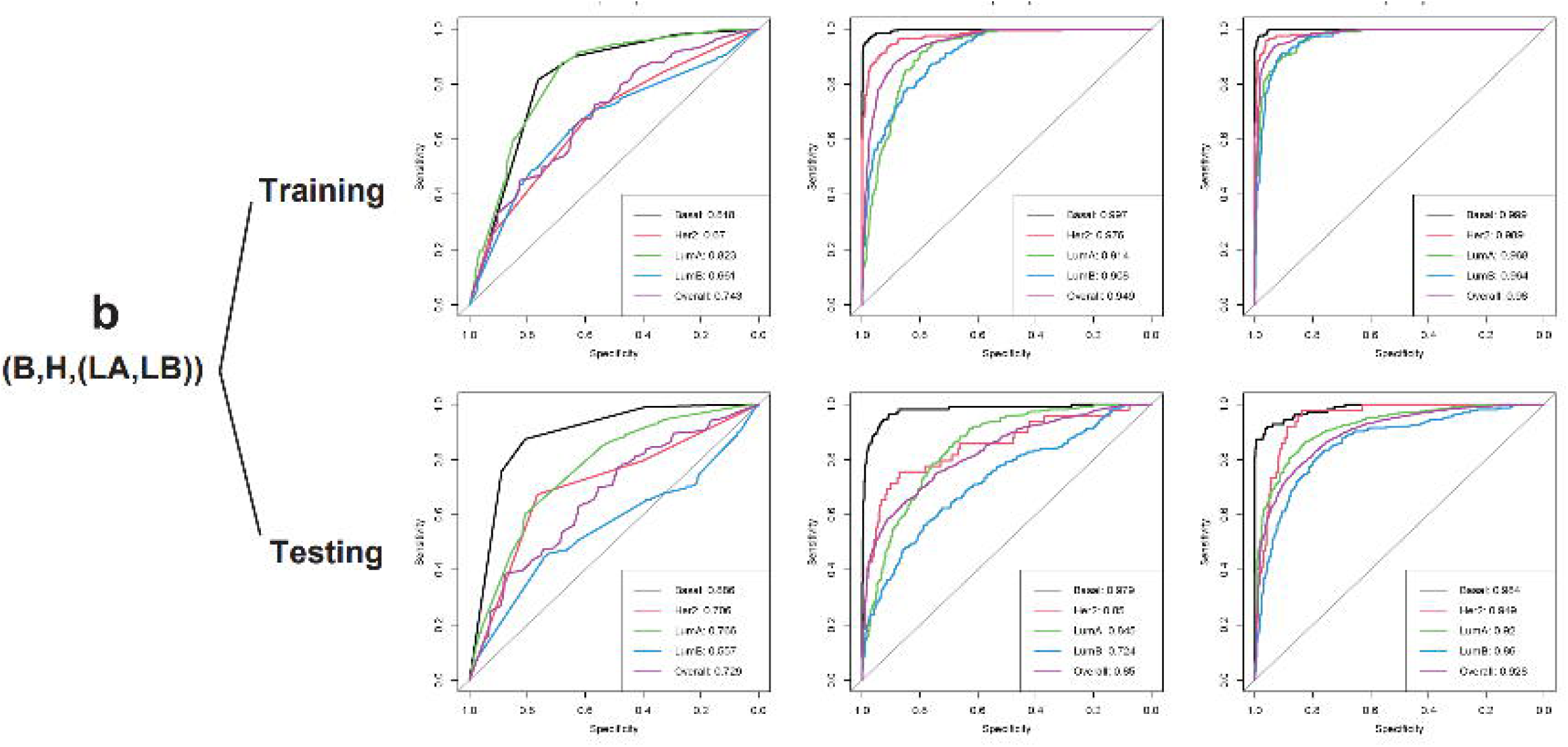
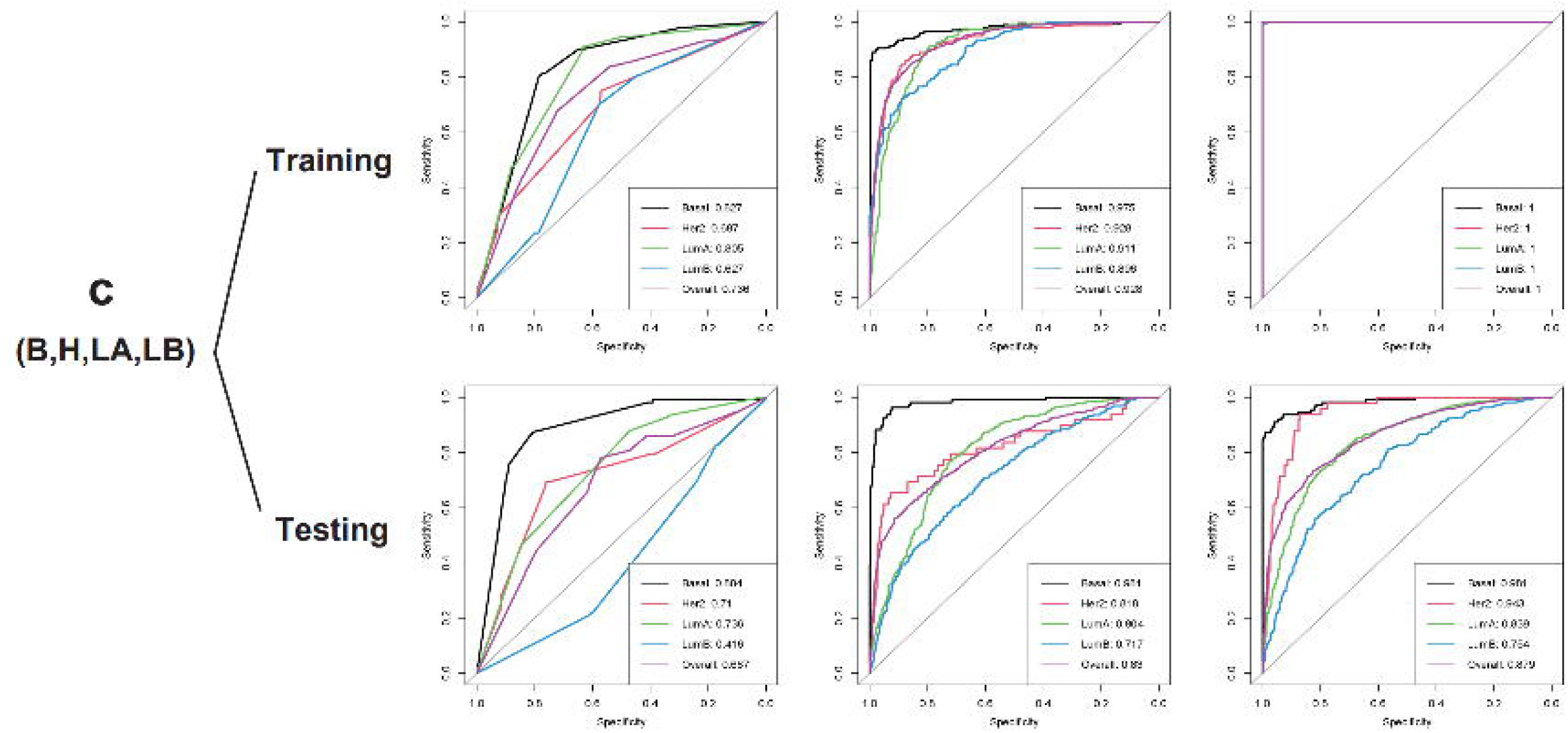

The heat patterns are in general consistent with the relative importance of DNA alteration classes previously discussed (**Fig. 4**). For examples, the B and (H,LA,LB) subtypes differed little in mutation profiles, but significantly in CNA and methylation profiles. According to the relative importance analysis, CNA and methylation alterations totally accounted for 87.02% of the feature importance for the *B-vs-(H,LA,LB)* sub-classifier. We also observed distinct signature heat patterns of respective DNA alteration classes, which are explained as follows:

First, gene mutation patterns showed substantial differences between the subtype groups with a total of nine signature mutated genes identified, including TP53, PIK3CA, GPR124, HRAS, SETD2, ERBB2, PNN, ATXN1 and CP (**Fig. 4a**). A few of these alterations were well-known cancer driving events. For examples, the TP53 gene, a tumor suppressor gene enabling tumors to be more aggressive, is typically mutated at the early stage of cancer to confer genetic instability to cancer cells and is associated with poor prognosis[63]. We observed that the TP53 gene was highly mutated in Basal and Her2 subtypes (**pattern ①**) and less mutated in Luminal-like subtypes. This suggested that patients with Basal and Her2 subtypes may have worse prognosis, as proven by many studies[1, 6, 7]. PIK3CA mutation was identified as (H,LA,LB) marker, which are most common in Luminal-like breast cancer, followed by Her2 breast cancer, and triple-negative breast cancer (**pattern ②**), consistent with previous studies[64, 65]. We also found HRAS, SETD2, ERBB2 gene mutations to be significantly associated with Her2 breast cancer, consistent with previous reports[66–68].

UGES also identified signature gene mutation as novel biomarkers of intrinsic subtypes, such as those affecting GPR124, PNN, ATXN1 and CP genes. Notably, mutations in the ATXN1 gene were newly identified by UGES to be associated with the LumB subtype. The major phenotypical difference between LumA and LumB subtypes is that LumB tumors express high level of Her2 and the proliferation marker Ki-67. Ki-67 is regulated by the Notch signalling pathway[69, 70]. Studies showed that ATXN1 is a negative regulator of the Notch pathway and ATXN1 knockdown enhanced tumor invasion by activating the Notch pathway. Therefore, a mutated ATXN1 gene may lose its inhibitor role of the Notch pathway and the abnormal expression of the Notch pathway leads to the overexpression of Ki-67 in Her2 cancer cells[71, 72]. GPR124 and CP gene mutations were also associated with tumor progression and invasion in breast cancer previously[73, 74], however not with specific intrinsic subtypes. While our results newly suggested these gene mutations are potential biomarkers of their specific intrinsic subtypes, more evidence is required to prove their functional roles.

Second, CNA alterations were found effective in identifying Basal and Her2 subtypes. As shown in **Fig. 4b** and **Table. 3**, Basal, Her2, LumA and LumB subtypes have distinct CNA patterns. For example, PGAP3 and MIR4728 gene copy numbers were highly amplified in the Her2 subtype (**pattern ③**), as compared to the Basal subtype. This is consistent with previous studies. For examples, PGAP3 might affect the Her2 subtype of breast cancer by being co-amplified with Her2[75]; MIR4728 gene is located within an intron of the ERBB2 gene, which is a well-known oncogene strongly associated with Her2 breast cancer[76]. Notably, UGES newly identified MIR5684 copy number deletion as a novel biomarker of Her2 subtype, while more evidence is required to prove the association.

Finally, in addition to genetic differences, epigenetic alterations such as methylation changes, also exhibited subtype-specific patterns (**Fig. 4c**). For example, as shown in **pattern ④**, nine genes (ADCY4, CRYAB, DNM3, HOXA11, INS, KCNH8, LAMB3, PRTN3, UCN) showed hypomethylation in the Basal subtype. While direct studies of these genes’ hypomethylation and the basal subtype were limited, hypomethylation of a gene generally elevates the gene’s expression level, and many of these genes were already shown to be highly expressed in triple negative breast cancer cells (TNBC), accounting for >80% of the Basal subtype. For examples, CRYAB was known to specifically enhance cell migration in TNBC cells[77]; and LAMB3 was known to mediate the apoptotic, proliferative, invasive, and metastatic abilities of Basal breast cancer cells[78]. UGES newly suggested AIRE and PRSS41 gene hypermethylation are also potential biomarkers of the Basal subtype breast cancer, which requires further experimental evidence.

## Discussion

In this study, we successfully constructed a DNA-level classifier UGES using only DNA alterations to identify breast cancer intrinsic subtypes. Along the development, we have gained several insights.

First, a multi-step strategy with an grouping of LumA and LumB subtypes is key to the better performance of UGES classifier. As the testing set result showed (**Fig. 2c**), the best AUC scores for single and overall subtypes were all obtained using multi-step strategies. (B,(H,(LA,LB))) had the best overall AUC scores, while (B,H,(LA,LB)) obtained the highest AUC scores 0.992 for identifying the Basal case. The AUC scores of the one-step (B,H,LA,LB) strategy were, however, the lowest consistently, in particular having trouble resolving the LumA and LumB subtypes, showing only AUC=0.852 and 0.782, respectively.

One potential explanation is that the one-step classifier may lose accuracy by ignoring the natural hierarchy of intrinsic subtypes. In fact, in the past, breast cancers have been broadly classified based on their gene expression profiles into Luminal- and Basal-type tumors. It is only recently that the Luminal type was further divided into two sub-groups[54]. This demonstrates the fact that Luminal-like subtypes differ more from Basal and Her2 subtypes as compared with each other. Thus, we conclude that the hierarchical stepwise classification method is a more realistic model in the identification of intrinsic subtypes. The inaccuracy of the PAM50 classifier is perhaps also attributable to the fact that it took a four-classification one-step approach.

Second, all types of DNA alterations are informative for defining subtypes. As shown in **Fig. 2d** and **Supplementary Fig. 2**, in the testing set among the four strategies, the overall AUC scores of single-feature classifiers were all over 0.6, with average AUC scores of 0.698, 0.854 and 0.919, respectively, for mutation, CNA and methylation features, suggesting that they are all informative. Notably, the AUC scores of methylation-only classifiers were comparable to the all-feature classifiers, the lowest of which was 0.879 in the one-step multi-classification strategy. In fact, methylation changes are becoming more and more accepted as biomarkers for breast cancer early detection[79]. Biologically, in the process of carcinogenesis, promoter hypermethylation is a more frequently occurring event than mutations[80], with estimates varying from 600 to 1000 aberrantly methylated genes per tumor[81]. This significant contribution of variability may explain the strong effect of methylation in defining subtypes and its high relative importance in our model.

Even while methylation as a whole has the main effect, mutations and CNAs remain a non-negligible change in individual signature alterations. For example, the feature alterations with top coefficient value in the *H-vs-(LA,LB)* sub-classifier is PGAP3 CNAs, and TP53 and PIK3CA mutations also ranked high in coefficient values in the *B-vs-(H,LA,LB)* and *H-vs-(LA,LB)* sub-classifiers, indicating that driving mutations and CNAs are still of vital importance in the intrinsic subtype classification of breast cancer. In recent years, many researchers used only methylation features to distinguish different breast cancer intrinsic subtypes[82–84], but these studies may not be sufficient according to our findings.

Another reason to favor an all-feature classifier over a methylation-only classifier is to avoid model overfitting. For example, the methylation-only (B,H,LA,LB) classifier performed better in identifying the Her2 subtype (AUC = 0.943) compared to the all-feature classifier (AUC = 0.935). However, if we looked into their specific training and testing performance, the overall AUC scores for the training and testing sets of the methylation-only (B,H,LA,LB) classifier were 1 and 0.879, while these for the all-feature (B,H,LA,LB) classifier, were 0.965 and 0.889, as shown in **Supplementary Fig. 2c**. The larger difference in training and testing performance of the methylation-only classifier demonstrated that it has high overfitting potential, and that only using methylation features to analyze breast cancer intrinsic subtypes would result in the loss of model generalizability. Thus, our analysis suggested constructing breast cancer intrinsic subtypes using all DNA alteration features in future studies.

Third, our findings can guide the further development of more specific and mutually exclusive DNA biomarkers aimed at distinguishing specific subtypes. For examples, among the 716 samples in which TP53 mutations occurred, only 199 had PIK3CA mutations; among the 824 samples with PIK3CA mutations, only 199 had TP53 mutations, demonstrating that PIK3CA mutations are highly mutually exclusive to TP53 mutations in breast cancer, which is consistent with previous studies[85].

It is worth to point out that our study has several limitations. First, we used PAM50 subtypes as initial training labels to construct the UGES classifier. In fact, the true intrinsic subtypes of breast cancer were extremely difficult to assess experimentally[4]. We used the PAM50 label because it is one of the most internationally accepted classification method for intrinsic subtypes. But instead of using expression data, we developed UGES using only DNA alterations, achieved highly accurate results, and produced clinically more distinguishable subtype groups, proving that the DNA alterations alone are predictive and reliable features. Second, public data for our analysis are still limited. While current combined sample size are adequate to clearly separate the major subtypes, minor subtypes were not identifiable because of the lack of training data. Researchers recently reported several refined intrinsic subtypes[86–88], which could be considered in future models as their data become more available. Third, so far, we did not consider chromosomal-level structural changes, which are fundamental for chromosomally instable breast cancers, and shall be considered in future models.

## Data Availability Statement

The multi-omics data underlying this article are public available in TCGA (https://gdc.cancer.gov/about-data/publications/pancanatlas) and two datasets in cBioPortal platform (http://www.cbioportal.org/). More details about the source code and the datasets for the analysis workflow presented in the article are given at https://github.com/labxscut/UGES.

## Author Contributions

JX, XL, YX and LCX conceived the study and designed the study. JX and LG collected the data. JX, BY and KL performed the modelling analysis. JX and WC conducted the interpretation of data. JX and LCX have drafted the work or substantively revised it. All authors read and approved the approved the submitted version.

## Acknowledgements

This study was funded by the Guangdong Basic and Applied Basic Research Foundation (2022A1515-011426 to LCX), National Natural Science Foundation of China (61873027 to LCX).

