## Supplementary_Table_1 for "Building a genetic and epigenetic predictive model of breast cancer intrinsic subtypes using large-scale data and hierarchical structure learning"

|  | IHC subtypes | | | |
| --- | --- | --- | --- | --- |
| PAM50 subtypes |  | Basal | Her2 | LumB |
|  | Basal | 110 | 12 | 17 |
|  | Her2 | 21 | 66 | 67 |
|  | LumA | 2 | 0 | 480 |
|  | LumB | 0 | 0 | 311 |
| Agreement | | 44.84% | | |

**Supplementary Table 1**. **PAM50 subtypes deviated substantially from IHC subtypes.** We can confirm Basal, Her2 and part of the Luminal B IHC subtypes in our data, among which is only 44.84% agreement with PAM50 subtypes.
