## Supplementary material for "Building a genetic and epigenetic predictive model of breast cancer intrinsic subtypes using large-scale data and hierarchical structure learning": Table_1

| **Table 1:** Clinical characteristics and multivariate cox regression results for the cohorts. | | | | | | | |
| --- | --- | --- | --- | --- | --- | --- | --- |
| **Data description** | | | | | | | |
| **Cohort** | | **Number of patients (missingness)** | | | **Ending events^1^** | | |
| TCGA | | 931 (5) | | | 122 | | |
| METABRIC | | 1134 (309) | | | 487 | | |
| Combined | | 2065 (314) | | | 609 | | |
| **Multivariate cox regression results** | | | | | | | |
| **Clinical characteristics** | | **Dataset** | **Count (%) / Mean (Range)** | **Hazard Ratio** | **95% CI** | **p-value** | **Significance^2^** |
| Age | At diagnosis | TCGA | 58.4 (26-90) | 1.038 | (1.024, 1.053) | 2.3e-07 | *** |
|  |  | METABRIC | 61.6 (21-96) | 1.041 | (1.033, 1.050) | < 2e-16 | *** |
|  |  | Combined | 60.1 (21-96) | 1.038 | (1.031, 1.045) | < 2e-16 | *** |
| ER status | Negative (Reference) | METABRIC | 218 (19.22%) |  |  |  |  |
|  | Positive |  | 916 (80.78%) | 0.749 | (0.563, 0.997) | 0.0474 | * |
| PR status | Negative (Reference) | METABRIC | 506 (44.62%) |  |  |  |  |
|  | Positive |  | 628 (55.38%) | 0.917 | (0.737, 1.140) | 0.4340 | n.s. |
| Her2 status | Negative (Reference) | METABRIC | 981 (86.51%) |  |  |  |  |
|  | Positive |  | 153 (13.49%) | 1.643 | (1.244, 2.169) | 0.0005 | *** |
| ^1^An ending event is defined as a patient death or data censor. ^2^Statistical significance is based on the fitted multivariate cox model. n.s.: not significant, *p < 0.05, **p < 0.01, ***p < 0.001. | | | | | | | |
