## Supplementary material for "Building a genetic and epigenetic predictive model of breast cancer intrinsic subtypes using large-scale data and hierarchical structure learning": Table_2

| **Table 2:** Relative importance of DNA alteration features in UGES sub-classifiers. | | |
| --- | --- | --- |
| **Sub-classifier** | **DNA alteration (# of)** | **Relative importance** |
| *B-vs-(H,LA,LB)*  *236 DNA features* | Mutation (48) | 12.68% |
|  | CNA (91) | 33.35% |
|  | **Methylation (97)** | **53.97%** |
| *H-vs-(LA,LB)*  *367 DNA features* | Mutation (105) | 19.78% |
|  | CNA (98) | 25.15% |
|  | **Methylation (164)** | **55.08%** |
| *LA-vs-LB*  *530 DNA features* | Mutation (227) | 28.63% |
|  | CNA (100) | 19.76% |
|  | **Methylation (203)** | **51.62%** |
