## Supplementary material for "Building a genetic and epigenetic predictive model of breast cancer intrinsic subtypes using large-scale data and hierarchical structure learning": Table_3

| **Table 3:** 52 signature subtype-delineating alterations for breast cancer that have been identified in differential analyses. | | | | | | |
| --- | --- | --- | --- | --- | --- | --- |
|  | ***B-vs-(H,LA,LB)*** | | ***H-vs-(LA,LB)*** | | ***LA-vs-LB*** | |
|  | **B marker** | **(H,LA,LB) marker** | **H marker** | **(LA,LB) marker** | **LA marker** | **LB marker** |
| **Mutation** | TP53 | PIK3CA | TP53  GPR124  HRAS  SETD2  ERBB2  PNN | / | / | ATXN1  CP |
| **CNA** | BCYRN1(+)  PGM3(+) | MIR4728(+)  MIR3677(+)  USP32(+) | PGAP3(+)  MIR5684(-) | SLC4A1(-)  PLAC8L1(+)  PSMC3IP(-)  SPRR2B(+) | MNX1(-)  DNAJC11(-)  ZNHIT6(-)  ZNF267(+)  C17orf105(-)  DHDDS(-) | KCNJ2(+)  FAM49B(+)  TMC8(+) |
| **Methylation** | AIRE(+)  PRSS41(+) | KCNH8(+)  DNM3(+)  UCN(+)  PRTN3(+) | CYBA(+) | LAMB3(+) | INS(+)  HOXA11(+)  KRT78(+)  PDZD4(+)  TNNT3(-)  PLEKHA6(+)  LINC00574(+)  NTF3(-) | PRDM14(+)  AQP1(+)  PENK(+)  CCDC65(+)  HTR1B(+)  ADCY4(+)  CRYAB(+) |
| Note: (+) denotes copy number amplification or hypermethylation, (-) denotes copy number deletion or hypomethylation. | | | | | | |
